# Chronic Aerobic Exercise Alleviates Amyloid-induced Capillary Dysfunction

**DOI:** 10.64898/2026.01.15.699787

**Authors:** Alfredo Cardenas-Rivera, Eda Erdogmus, Austin Birmingham, Evan Yee, Jaime Anton, Zhengyi Lu, Yuntao Li, Neha Singh, Matthew Morais Johnson, Nazila Loghmani, Chang Liu, Mohammad A. Yaseen

## Abstract

Age-related cerebrovascular dysfunction is increasingly recognized as a critical contributor to cognitive decline and Alzheimer’s disease (AD) progression. Aerobic physical activity (PA) and other modifiable lifestyle interventions can substantially reduce the likelihood of dementia; however, their ability to mitigate cerebrovascular alterations remains poorly defined. PA reportedly improves systemic vascular health and cognitive function in aging humans, but its impact on cerebrovascular function during aging and amyloid β (Aβ) pathology is unclear. Here, we longitudinally quantified microvascular oxygen tension and stimulus-evoked oxygen dynamics in awake APP/PS1dE9 mice and wild-type littermates using two-photon phosphorescence lifetime microscopy. Routine aerobic PA initiated in early adulthood preserved basal arteriolar, capillary, and venular oxygenation, prevented age-dependent increases in microvascular heterogeneity, and mitigated excessive oxygen extraction in preclinical AD mice. While amyloid pathology impaired stimulus-evoked oxygen responses across vascular compartments, PA selectively enhanced capillary dilation and accelerated hyperemic kinetics without altering vascular density or architecture. Notably, sedentary AD mice developed lower, widely-dispersed distributions in capillary oxygenation, hallmarks of malignant microvascular dysfunction, which were largely absent in physically active animals. These findings demonstrate that routine aerobic PA preserves basal capillary oxygenation and stimulus-evoked hyperemia during aging and Aβ, supporting a capillary-centric mechanism through which exercise confers neurovascular resilience in preclinical AD.

## 1. INTRODUCTION

Amyloid-β (Aβ) accumulation and misfolded tau protein are traditionally considered the most distinctive attributes of Alzheimer’s disease (AD) pathology [1–5]. However, mounting evidence indicates that cerebrovascular dysfunction precedes and accelerates neurodegeneration and plays an integral role in preclinical AD progression [6–10]. Alterations in cerebral blood flow, oxygen extraction, and neurovascular coupling are detectable in cognitively normal individuals at risk for AD [11–16] and are strongly associated with early cognitive impairment [10,17–19]. Functional neuroimaging studies consistently report aberrant blood-oxygen-level-dependent (BOLD) responses during aging and preclinical AD [20–22], yet the microvascular mechanisms underlying these changes remain inadequately understood.

The cerebral microvasculature, particularly the capillary network, plays a central role in oxygen delivery, metabolite exchange, and clearance of microenvironmental toxins, including Aβ [23,24]. Emerging work suggests that capillary dysfunction can generate localized hypoxia and metabolic stress even in the absence of overt vascular rarefaction [25,26]. Such disruptions, including impaired perfusion, increased flow heterogeneity, and stalled red blood cells, are not uniformly distributed, and consequently, they create microscopic territories of impaired oxygen delivery, a phenomenon increasingly implicated in both normal aging and AD pathogenesis [27]. Despite this, most studies of cerebrovascular aging rely on bulk measures of blood flow or oxygenation, obscuring capillary-scale dysfunction that may critically influence brain tissue’s heterogeneous vulnerability.

The impacts of modifiable risk factors on AD progression and other dementias continue to grain recognition as promising preventive measures. Reports estimate that between 30-50% of dementia cases can be prevented by treating obesity, inadequate sleep, hypertension, physical inactivity, and other prevalent health risks [28–32]. Among those factors, aerobic physical activity (PA) is one of the most accessible and robust non-pharmacological interventions known to preserve cognitive function. In aging humans, 3-12 months of moderate PA has been shown to improve executive function, memory, and processing speed, while reducing the risk of dementia [33–36]. At the vascular level, improvements in endothelial function, arterial compliance, and systemic oxygen delivery have been observed during acute PA [35,37–42]. In the brain, exercise has been shown to improve cerebral blood flow, reduce neuroinflammation, basal oxygenation, and attenuate amyloid burden in animal models of normal aging and models of AD [43–50]. However, a detailed understanding of chronic PA’s long-term impacts on microvascular oxygen delivery during aging coupled with AD progression remains incomplete.

Critically, neurovascular coupling—the process by which neuronal activity drives local increases in blood flow and oxygen delivery—relies on coordinated signaling among endothelial cells, vascular smooth muscle cells, and pericytes [13,51–53]. These cellular elements are particularly vulnerable to Aβ, which disrupts vasomotor control, capillary dilation, and oxygen extraction efficiency [10,25,54–60]. At the same time, PA is known to modulate the phenotype and function of these vascular cells, suggesting that exercise may exert its neuroprotective effects, at least in part, through restoration of microvascular dynamics rather than gross structural remodeling.

Here, we test the hypothesis that chronic aerobic PA preserves microvascular oxygenation under basal conditions and during functional hyperemia during aging and amyloid pathology. Using longitudinal two-photon phosphorescence lifetime microscopy in awake mice, we directly measured intravascular oxygen tension, oxygen extraction fraction, and stimulus-evoked oxygen responses across arterioles, capillaries, and venules in APP/PS1dE9 mice and wild-type controls. By combining high-resolution oxygen measurements with functional stimulation and vascular morphology analysis, we delineate how PA modifies capillary oxygen delivery, heterogeneity, and responsiveness independently of vascular density. Our findings identify capillary oxygen dysfunction as a key target of PA and provide mechanistic insight into how exercise may counteract early neurovascular impairments in preclinical AD.

## 3. RESULTS

Physical exercise reportedly enhances systemic vascular function and memory in healthy adult humans [61]. To determine whether physical activity mitigates cerebrovascular alterations during aging and preclinical AD, we longitudinally measured microvascular oxygenation in the somatosensory cortices of awake, head-restrained mice. Male APP/PS1dE9 transgenic (AD) mice and wild-type littermates (WT) were categorized into 4 cohorts. From age 6-12 months, 2 cohorts (AD-PA and WT-PA) underwent structured treadmill running exercise (30 minutes/day, 5 days/week; Fig. 1A), while the remaining cohorts remained sedentary (AD-noPA and WT-noPA). Using two-photon phosphorescence lifetime microscopy and the phosphorescent oxygen sensitizer Oxyphor2P, intravascular oxygen tension (pO2) was quantified up to 375 μm below the cortical surface in response to somatosensory whisker stimulation after 1 month (phase 1) and 6 months (phase 2). The influences of Aβ pathology, PA, and age were evaluated using linear mixed-effect modelling.

**Figure 1.**
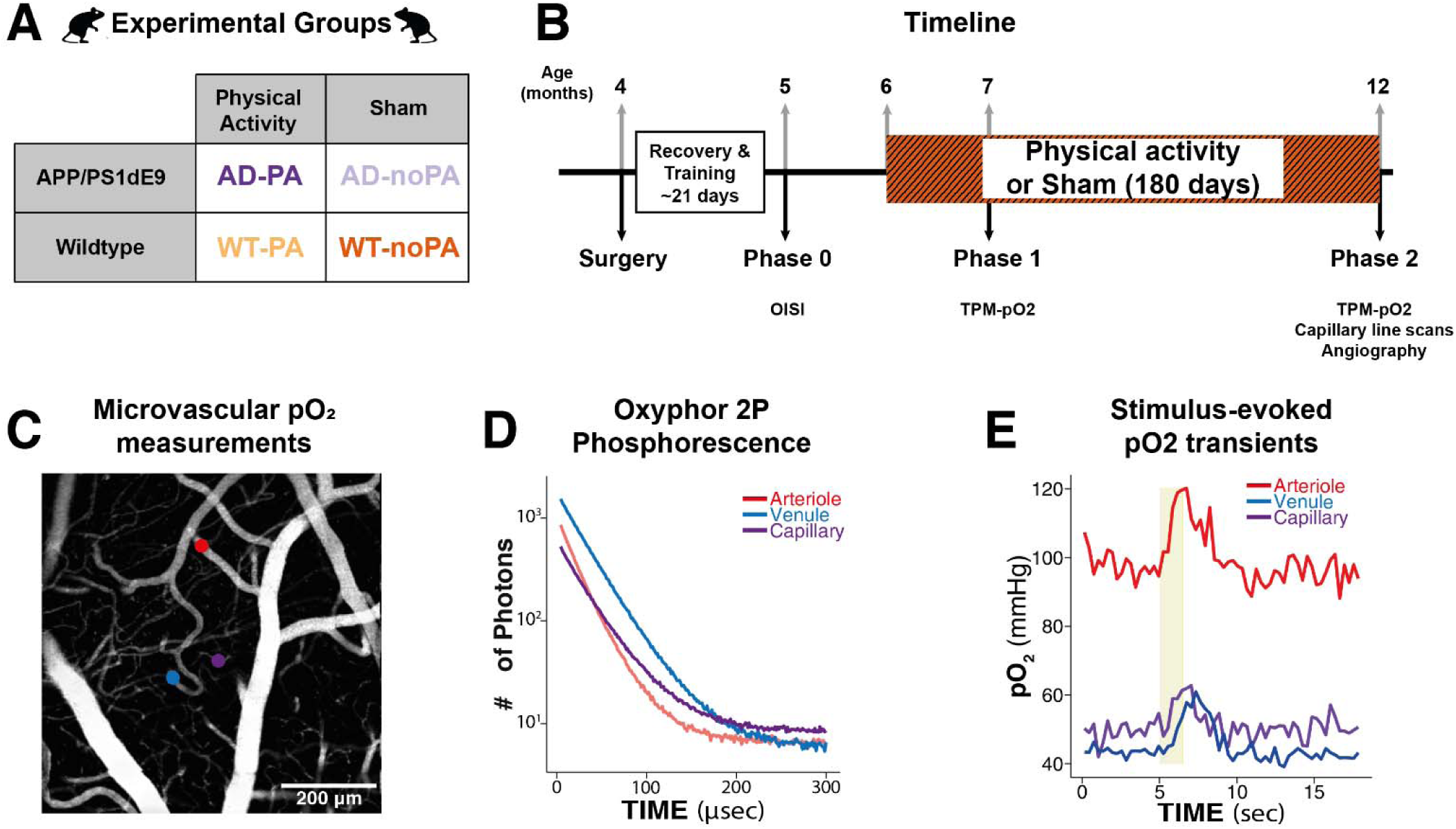
Experimental study details. 4 mouse cohorts (n = 6 each, male) underwent chronic cranial window implantation at age 4 months. 2 cohorts (AD-PA and WT-PA) underwent routine treadmill exercise (11/m/min, 30 minutes/day 3 days/week) from age 6-12 months while the other 2 cohorts remained sedentary. At Phases 1 and 2, microvascular pO2 was measured at different cortical depths in arterioles, capillaries, and venules via phosphorescence lifetime imaging of the oxygen sensitizer Oxhyphor2P. Microvascular angiography and line scan measurements of capillary diameter were collected at Phase 2.

### 3a Basal oxygenation

Figure 2 displays baseline intravascular pO2 from cortical arterioles, capillaries, and venules of each experimental cohort. After one month (Phase 1), both AD and WT mice undergoing PA demonstrated significantly higher levels of arteriolar oxygen compared to their sedentary counterparts (Fig. 1A). Significant differences in arteriolar pO2 were sustained between PA and sedentary cohorts after 6 months (phase 2); however, arterial pO2 also declined with age in both AD mouse cohorts. In WT animals, arteriolar pO2 at phase 2 was significantly higher for the WT-PA animals relative to WT-noPA.

**Figure 2.**
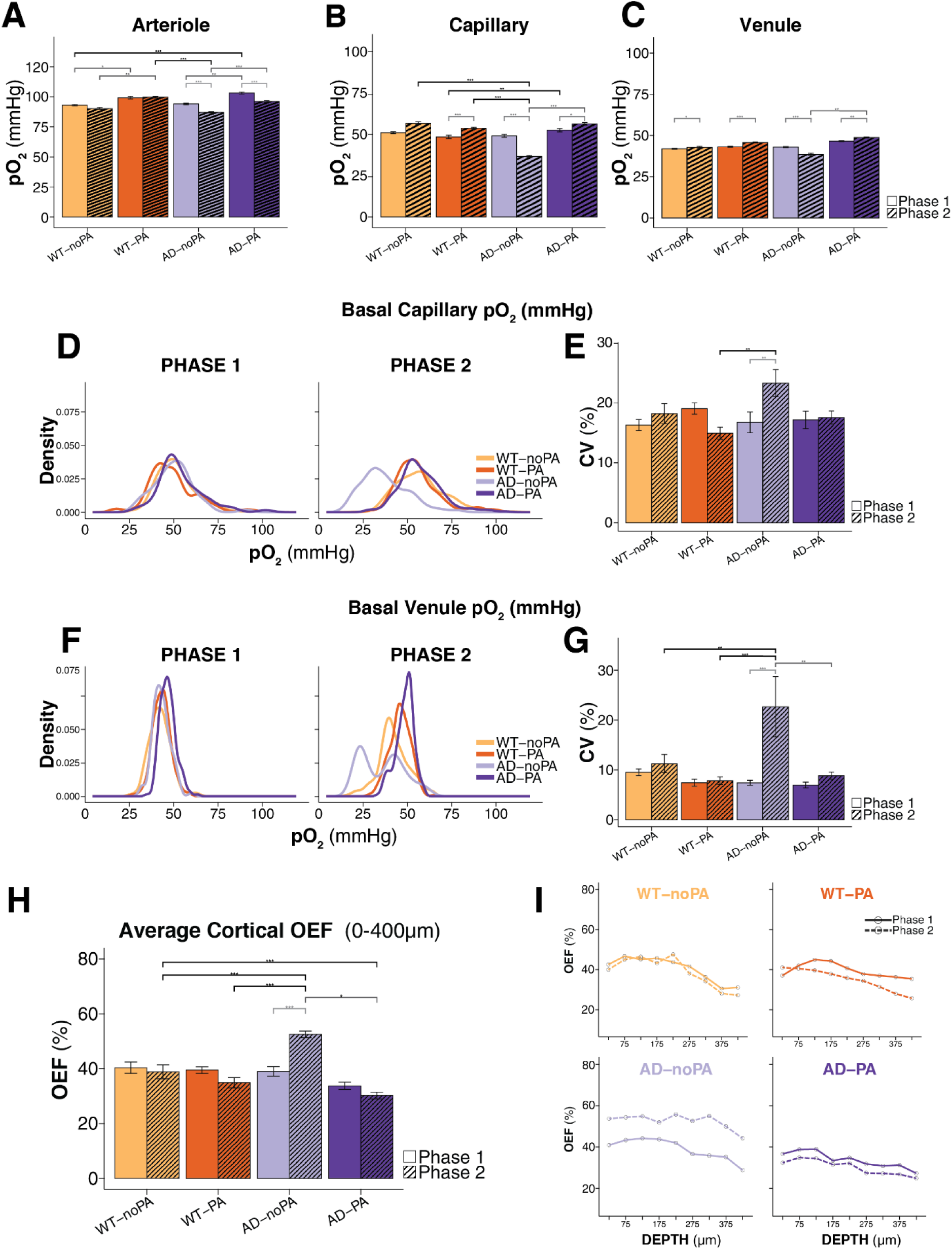
Basal microvascular pO_2_ in AD and wildtype mice. ***(A)*** Arteriolar cortical oxygenation remains stable between 7 months (phase 1) and 12 months (phase 2) in wild-type (WT) mice, but is significantly reduced in 12-month-old AD mice. Physical activity (PA) induces an early and sustained increase in arteriolar oxygenation. ***(B)*** Capillary oxygenation increases with aging in WT mice and is markedly reduced in AD-noPA mice at 12 months; this reduction is prevented by PA. ***(C)*** Venular oxygenation remains stable across ages and groups, except for a significant decline in AD-noPA mice at 12 months. ***(D-G)*** Basal pO_2_ distributions at the capillary and venular levels are similar for all groups at 7 months. AD-noPA mice at 12 months display a left-shifted, bimodal distribution, while other groups maintain a narrow, unimodal profile. ***(H-I)*** The cortical oxygen extraction fraction (OEF) increases for AD-noPA mice between phases 1 and 2. Routine PA yielded lower cortical OEFs in WT and AD mice at phase 2.

At the capillary level, where the majority of metabolite exchange and Aβ clearance reportedly transpires [62], PA yielded pronounced differences in AD mice at Phase 2. Both WT-PA and AD-PA animals had significantly higher baseline capillary oxygenation at phase 2 relative to phase 1. Conversely, AD-noPA mice demonstrated a pronounced decline in capillary oxygenation over 6 months, and capillary pO2 did not change significantly in WT-noPA mice. Similar trends were observed in the venous compartment. In AD-noPA mice, baseline venular oxygenation decreased significantly. In all other cohorts, venous pO2 increased significantly between phases 1 and 2.

Microvascular heterogeneity has recently emerged as an important indicator of pathological disturbances, reflecting microscopic territories of brain tissue with disrupted metabolic supply [63,64]. The distribution profiles and coefficients of variations (C.V.) of baseline intravascular pO2 in Figs 1.D-G highlight how Aβ pathology, age, and PA alter microvascular heterogeneity through their impacts on dispersity of capillary and venous oxygenation. In all groups during Phase 1, the distributions of capillary oxygen tension exhibited a unimodal Gaussian profile centered around ∼50 mmHg with similar C.V. at 7 months. By 12 months, capillary pO2 within sedentary AD mice differed substantially from the other cohorts, displaying a lower, bimodal distribution. The broader distribution of reduced pO2 is consistent with widespread, heterogeneous capillary hypoxia. Venular oxygen distributions paralleled these trends, remaining unimodal (∼45 mmHg) at 7 months and becoming bimodal in sedentary AD mice at 12 months, with distinct modes at ∼23 mmHg and ∼45 mmHg. In AD-PA mice, PA mitigated increases in malignant capillary and venous heterogeneity, maintaining values comparable to those observed in age-matched WT controls (Fig. 2E,G).

We calculated oxygen extraction fraction 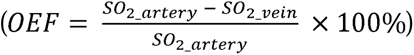 to quantify the amount of oxygen delivered at each cortical depth. Microvascular pO2 measurements were converted to oxygen saturation using the Hill equation [65]. Figs 2.H-I depict average and depth-resolved OEF in each cohort at each phase. At phase 1, all groups demonstrated OEFs ranging around 40% that decreased slightly with cortical depth, in agreement with prior studies of 3-6 month old awake mice [66–68]. Between phases 1 and 2, AD-noPA animals experienced significant increases in OEF across all layers, while all other cohorts showed slight reductions. At phase 2, OEF in animals with PA appeared lower than their sedentary counterparts.

### 3.b Functional Hyperemia

We applied intravital Optical Intrinsic Signal Imaging (OISI) and 2-photon phosphorescent lifetime imaging 2P-PLIM to assess stimulus-evoked hyperemic responses in microvascular pO_2_ and the impacts of physical activity PA, Aβ, and age. We measured transient changes in cortical blood volume and intravascular pO2 in response to 3-second pneumatic whisker deflection. OISI using 532 nm reflectance was first applied to identify the activated cortical region. Stimulation trials were then conducted to measure pO2 in individual microvessels from the cortical surface down to 350 μm in 50 μm increments. Linear mixed effect modeling revealed no significant depth-related differences. As a result, data from cortical layers were combined.

Figs. 3A-C displays OISI-based measurements of cerebral blood volume changes after 0 months (baseline), 1 month (phase 1), and 6-months of aerobic PA. Reductions in 532 nm OISI reflectance intensity correspond to increases in HbT, allowing simple quantification of cerebral blood volume [69]. Linear mixed effects analysis revealed no significant differences between experimental groups or phases for hyperemic blood volume changes. Figs 3D-E highlight group-averaged capillary pO_2_ transients at phases 1 and 2, obtained by 2P-PLIM. In all measurements, capillary pO2 reached maximal levels shortly after stimulus cessation, slower than peak changes in blood volume. In WT animals, peak amplitude of capillary pO2 remained unaffected by age or PA. Conversely, both AD-PA and AD-noPA mice demonstrated markedly lower peak responses at 12 months. Figs 3 G-H display maximum stimulus-induced pO2 in arterioles, capillaries, and venules, resolved into baseline pO2 and stimulus-induced hyperemic increases (colored bars and white bars, respectively). Statistical bars in figs 3. G-H indicate significant differences in maximum pO2 during stimulation. In sedentary AD mice, maximal pO2 were significantly lower in older animals (phase 2) within all vascular compartments. While in both sedentary and active WT animals, peak pO2 either increased (capillaries and venules) or remained unchanged. Peak pO₂ remained stable across time points in WT animals, with no differences between sedentary and active groups.

**Figure 3.**
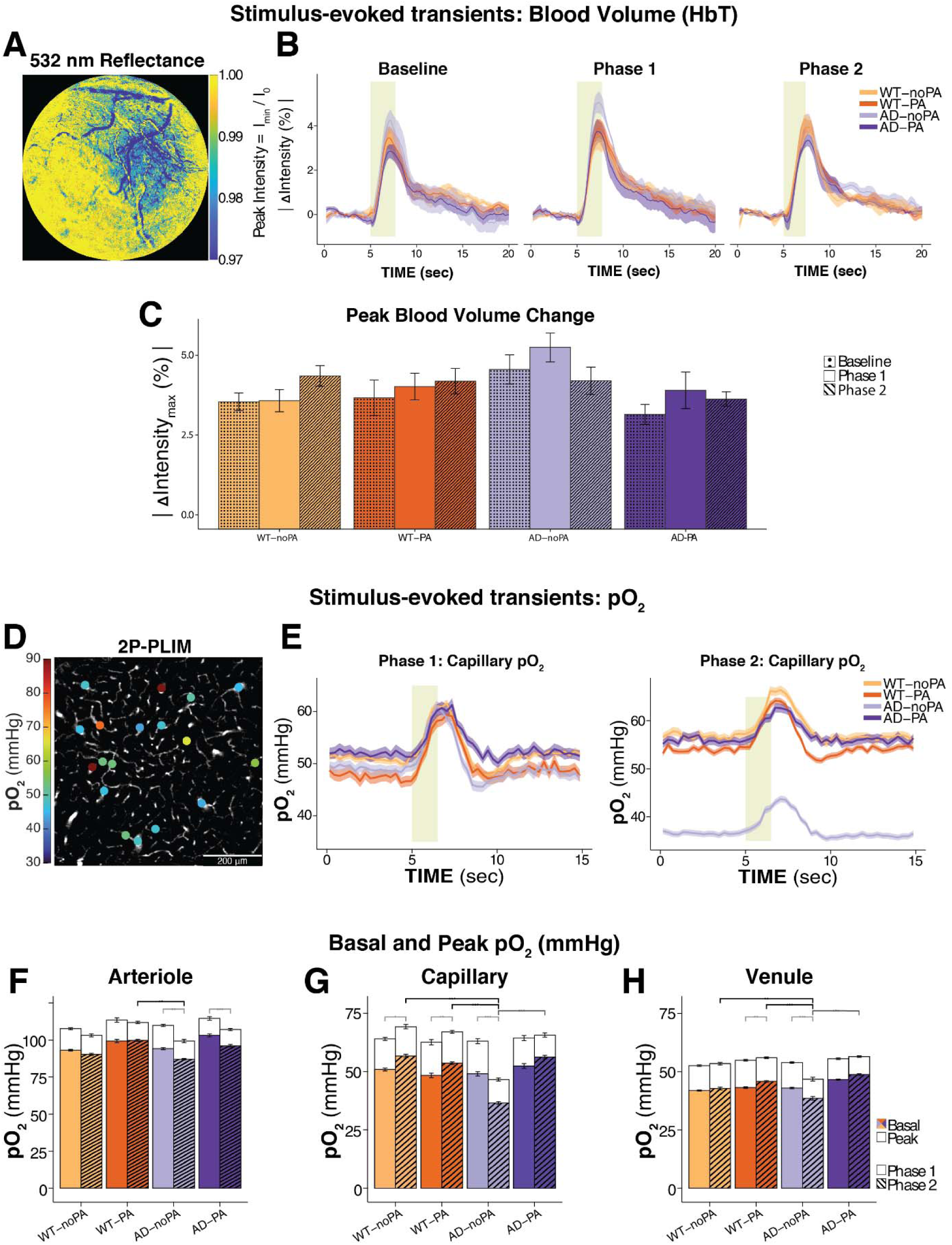
Hyperemia dynamics during functional stimulation. (**A-C**) Representative OISI image and group-averaged transients of 532 nm reflectance intensity. Pneumatic whisker stimulation provoked localized increases in blood volume within the somatosensory cortex. Hyperemic blood volume changes did not vary significantly between experimental groups or phases ***(D-E)*** Pointwise 2P-PLIM measurements in cortical microvessels. 6 months of routine aerobic PA protects AD mice from reductions in capillary pO2 ***(F-H)*** Alterations in basal and peak pO2 in arterioles, capillaries, and venules. Statistical bars in figs f-h indicate significant differences in maximum pO2 during stimulus trials. Peak differences in arteriolar oxygenation are higher in sedentary groups of both genotypes. In capillaries, AD-noPA mice display lower baseline pO2 but larger ΔpO2 than AD-PA at both 12 months. Venular oxygenation peaks increased slightly between phases 1 and 2 in WT-PA mice. PA prevented reductions in venous oxygenation in AD-mice.

To comprehensively assess variations in total blood oxygen, vascular pO2 was converted to oxygen saturation (SO2) using the Hill equation [65,67]. Fig 3 displays group-averaged SO2 transients for each vascular compartment and the calculated metrics for stimulus-evoked hyperemia (ΔSO2 = SO2_peak_ ^-^ SO2_baseline,_ AUC = area under curve). In arterioles, baseline SO2 levels at phase 2 were slightly higher for WT-PA and AD-PA animals relative to the noPA groups. In all groups except AD-PA, functional stimulation evoked smaller increases in vascular oxygenation at phase 2, as indicated by ΔSO2 and AUC plots (figs 4.a and b). Corresponding profiles for pO2 are displayed in supplementary fig.1

Perivascular amyloid-β accumulation reportedly induces capillary endothelial dysfunction and impairs amyloid clearance, establishing a feed-forward degenerative loop [70]. Notably, our observations indicate that PA attenuates endothelial dysfunction and limits this pathological amplification. In WT animals, stimulus-evoked capillary SO2 changes were lower at phase 2 relative to phase 1, and WT-PA animals showed slightly higher SO2 changes compared to WT-noPA animals (Fig 4C). While AD-PA animals maintained higher baseline SO2 levels, stimulus-evoked changes in SO2 were smaller in AD-PA animals and reduced further between phases 1 and 2 (Fig 4D). Interestingly, hyperemic responses in AD-PA animals displayed faster kinetics compared to the other groups, evidenced by lower rise times and fall times (Fig 4D). Both WT-PA and AD-PA animals showed slightly faster responses than their sedentary counterparts. These differences were more pronounced at phase 2 compared to phase 1. The results strongly indicate that PA- and aging induce separate, and counteracting variations in pericyte-mediated capillary constriction [71,72].

**Figure 4.**
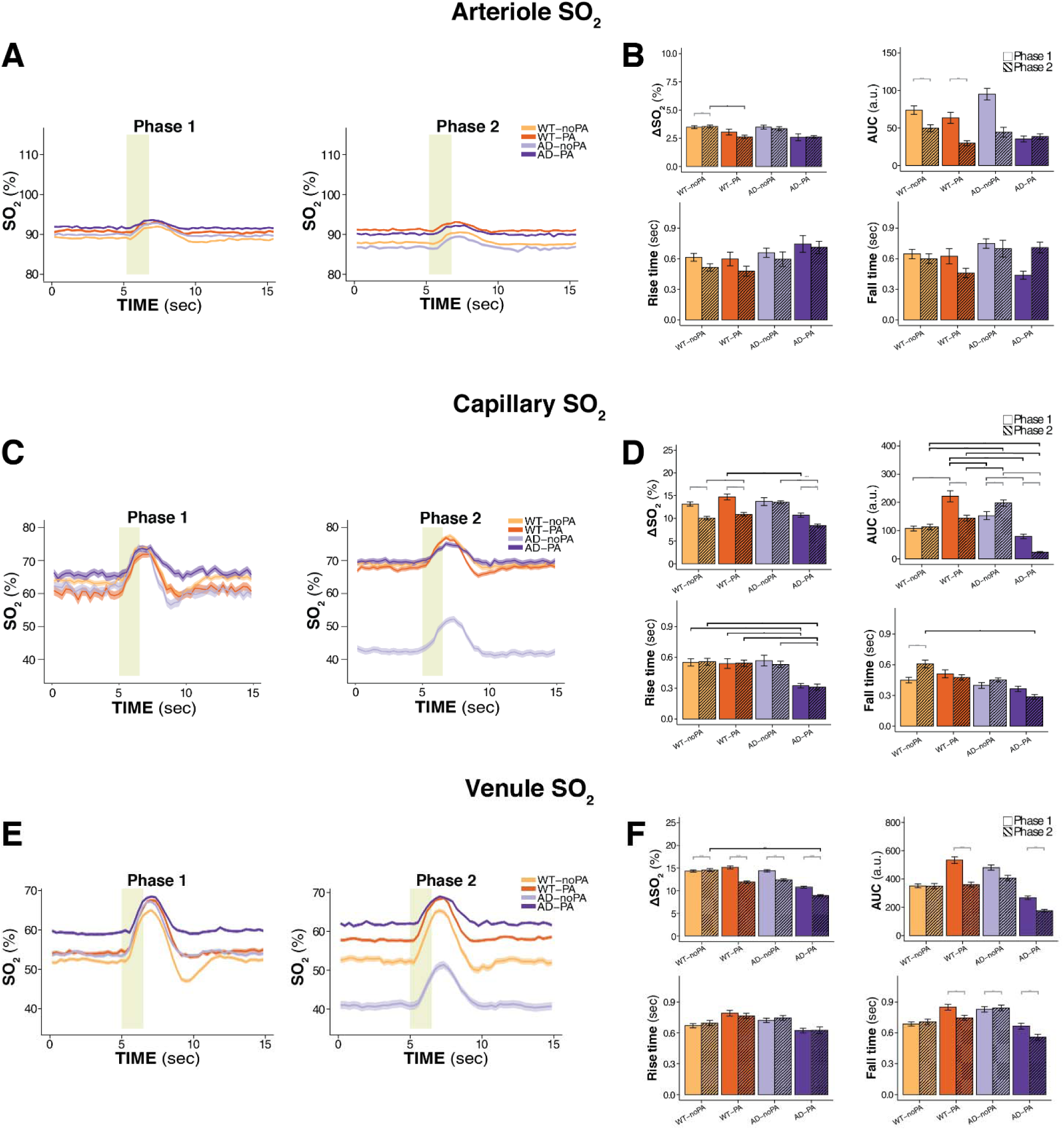
Microvascular SO_2_ transients during whisker stimulation in wild-type and AD mice under aging and physical activity: **(A-B)** Arteriolar oxygen transients show higher basal SO_2_, but smaller stimulus-evoked increases, in both AD-PA and WT-PA groups. For all groups, functional stimulation evoked smaller arteriolar SO_2_ changes at phase 2 than phase 1. **(C-D)** In capillary vessels, Aβ and aging yielded reductions in basal SO_2_ in AD-noPA mice. Routine PA protected AD-PA mice from reductions in basal SO_2_. In WT animals, hyperemic increases in SO2 were higher in PA animals compared to sedentary noPA animals, while in AD animals, PA animals responded to stimulation with smaller, yet faster changes in SO2. (**E-F**) In venules, differences in basal SO2 were higher in WT-PA and AD-PA animals at phase 2. Functional stimulation evoked smaller venous SO_2_ changes at phase 2 than phase 1.

In venules, observations of baseline and stimulus-evoked SO2 responses appeared to follow those of the capillary vessels (figs 4.e-f). AD-noPA animals showed lower baseline SO2 compared to AD-PA animals, and these differences were more pronounced at phase 2. Stimulus-evoked hyperemic SO2 changes were lower at phase 2 relative to phase 1. Interestingly in both PA groups, we observed age-related variations in fall time of the SO2 response. Both PA animals showed faster fall times at phase 2 compared to phase 1.

At Phase 2, after 6-months of PA, we applied rapid, high-speed line-scanning to monitor capillary diameter during whisker stimulation trials (Fig. 5A). In the activated cortical region, we observed two distinct populations of capillaries. One set demonstrated stimulus-evoked vasodilation in response to pneumatic air puffs (“responsive” capillaries; Fig. 5C), while the other set of capillaries exhibited no dilatory response (“non-responsive” capillaries; Fig. 5B). Fig 5D display resting, pre-stimulus diameters for both capillary populations, along with their relative proportions for each cohort at phase 2 (fig 5E). For all four experimental cohorts, no appreciable differences were detected between responsive and non-responsive capillary populations for resting state diameters and their relative proportions. In all cases, average capillaries diameters remained tightly distributed around 4.9 µm. Fig 5F displays peak stimulus-induced dilation of responsive capillaries in both WT and AD mice. In both cohorts, mice that underwent six months of PA demonstrated significantly larger peak dilation relative to their sedentary counterparts. Intriguingly, responsive capillaries in AD mice demonstrated larger dilations compared to WT mice, despite having similar baseline capillary diameters. The maximal dilation occurred in response to the first air-puff across all experimental groups, while the maximal oxygenation peak occurred following the final air-puff (Fig. 5D). The area under the curve (AUC) of capillary diameter change indicates a greater overall vascular response in PA mice, compared to sedentary animals (Fig. 5G). AD mice exhibited larger AUC values than WT counterparts, with the most pronounced response observed in the AD-PA group. Fig 5H quantifies the heterogeneity in peak dilation among responsive capillaries. AD mice showed higher, yet not significantly different, heterogeneity in stimulus-evoked diameter changes.

**Figure 5.**
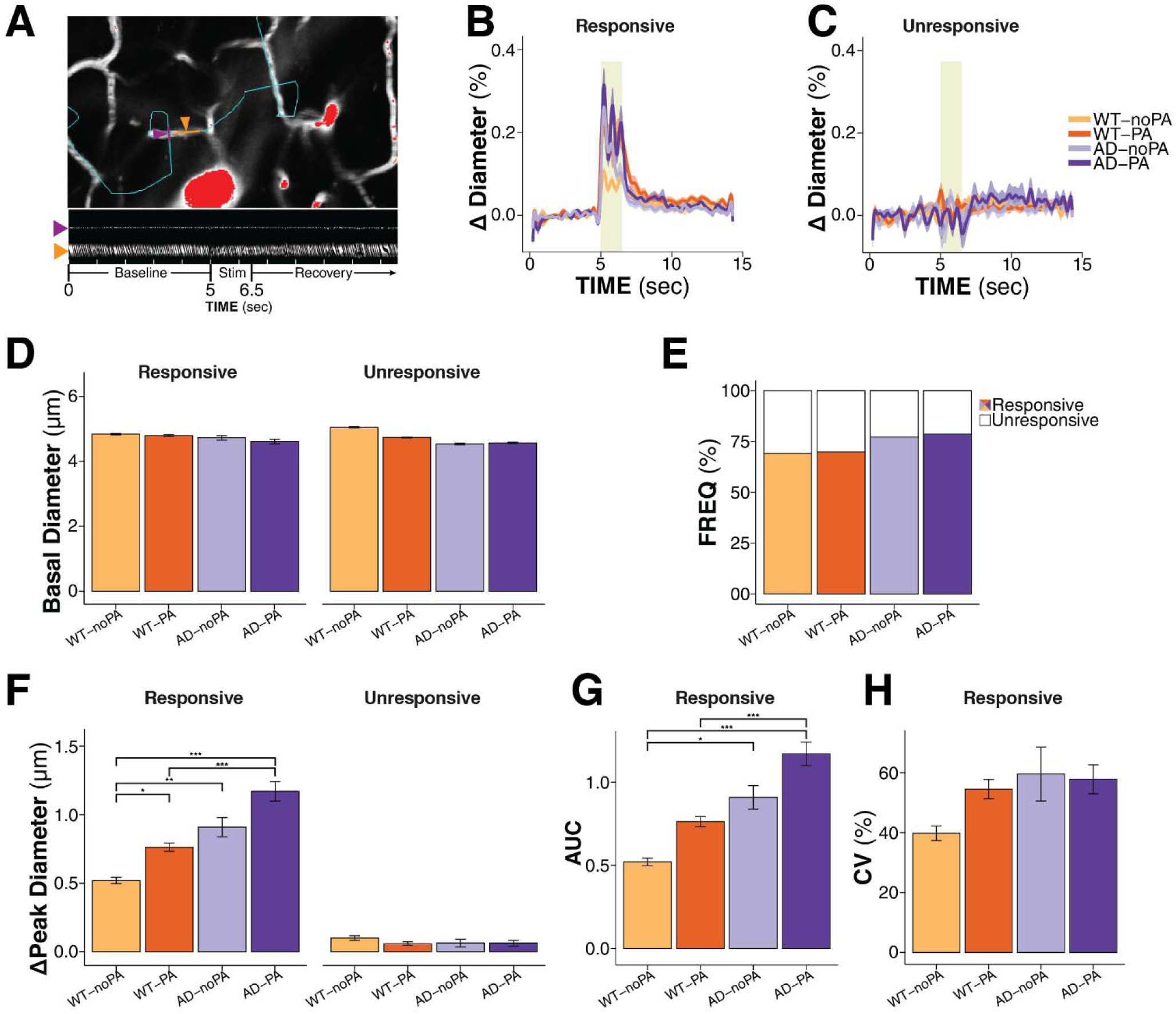
Capillary dilation responses during functional activation in wild-type and AD mice at Phase 2 (age 12 months). **(A)** Line-scan imaging protocol used to measure rapid vascular dynamics during whisker stimulation. **(B–C)** Two distinct subtypes of capillary responses were identified: responsive vessels exhibiting rapid, stimulus-locked dilations to each pneumatic air-puff and (**C**) non-responsive vessels displaying no discernible change in diameter. **(D, E**) Baseline diameters for responsive and non-responsive capillaries at phase 2. **(F-G**) Quantitative analysis of stimulus-evoked capillary dilation, area under the curve (AUC), and heterogeneity revealed larger vasodilatory responses in trained AD-PA and WT-PA mice after 6 months of regular physical activity **(H**) computed coefficients of variation from responsive capillaries indicate no differences in heterogeneity of capillary dilation.

### 3.c Microvascular morphology

At phase 2, we quantified features of microvascular angiography to assess the influences of Aβ pathology and 6 months routine PA. High-resolution z-stacks of cortical microvessels (∼450 μm thick at 1 µm intervals) were acquired using Dextran-labeled fluorescein. We performed segmentation and analysis of the vascular image stacks using Matlab-based VIDA software package [73]. Supplementary fig 2 displays example stack images for each cohort and computed morphological metrics for the vascular stacks. Although linear mixed effect modeling identified differences at specific cortical depths between some experimental groups, we observed no discernible trends in vascular density, vessel length, cross sectional area of microvessels, or total vascular surface between sedentary and trained groups, irrespective of genotype. These results indicate that neither Aβ accumulation nor chronic exercise appreciably alter microvascular architecture in the barrel cortex at age 12 months.

## DISCUSSION

Numerous studies have demonstrated that routine engagement in aerobic (PA) helps maintain and improve cognition and memory in aging humans and animals. In particular PA, offers benefits for processing speed, attention, and executive control [74,75]. However, a complete understanding of PA’s effects on cellular and vascular physiology in the brain remains unclear. Our findings provide compelling evidence that routine aerobic PA mitigates Aβ’s detrimental effects on cortical oxygen supply, oxygen extraction fraction, vascular heterogeneity, and vascular responses to functional stimulation. These results offer novel insights into PA’s neurovascular implications highlighted in previous studies in healthy humans and AD patients [61,76–78]. Specifically, our findings indicate that both AD pathology and PA do not affect the vascular network uniformly. The results offer greater insight regarding variations between healthy aging and AD progression observed via fMRI BOLD neuroimaging.

We observed age-associated reductions in basal oxygen tension across the cortical vascular network in sedentary APP/PS1dE9 mice (AD-noPA), extending from arteriolar inflow to venular outflow. Though the mechanism remains unclear, the persistent reduction in arteriolar pO is consistent with earlier reports that attribute vascular abnormalities to cerebral amyloid angiopathy (CAA) along cerebral arteries. In the APP/PS1dE9 mouse model, CAA begins to manifest around six months of age and has been shown to promote multiple impairments to arterial structure and function, including degeneration of vascular smooth muscle cells (vSMCs), compromised integrity of the periarterial basement membrane, impaired vessel wall compliance, and vasomotor function [79]. Our basal pO2 observations in AD-noPA mice demonstrated more substantial alterations in the capillary and venous compartments at age 12 months (phase 2). In addition to significant age-related reductions in basal pO_2_, the observed increases in heterogeneity of capillary and venous pO_2_ reflect variable, uneven perfusion and functional shunting within the capillary beds. Such alterations purportedly result in microscopic territories of hypoxic tissue as blood flow and oxygen availability drops [63,64,80]. We observed no depth-dependent differences; however, our OEF observations agree well with prior 2P-PLIM observations in awake female C57BL/6, 3–5 months mice [66,67]. In our AD-noPA mice, OEF increased substantially between ages 7 months (phase 1) and 12 months (phase 2). Conversely, based on bolus tracking measurements and Jespersen & Østergaard’s 3-compartment model, Gutiérrez-Jimenez et al computed lower OEF values in anesthetized APP/PS1dE9 mice and wildtype littermates at age 18-months [63,81]. The OEF disparities are likely attributable to different measurement methods, anesthesia-induced alterations to cerebral energy metabolism and blood flow, and age-related variations in cerebral blood flow. When taken together, however, the findings from our study and Gutiérrez-Jimenez highlight how capillary function and perfusion each vary independently with age and during AD progression. The resultant age-dependent alterations in cerebral blood flow, metabolism, and oxygen extraction likely account for differences in fMRI BOLD response for pre-symptomatic, subclinical, and demented AD patients [80].

Routine aerobic PA yielded significant increases in basal arteriolar pO2 in both WT and AD mice at phases 1 and 2. In capillaries and venules, PA prevented or mitigated reductions in basal pO2 in AD mice at phase 2. We observed that 6-months of routine, forced PA mitigated AD-related reductions to basal arteriolar pO2 and AD-related increases to basal OEF, in agreement with prior studies that assessed voluntary PA in younger APP/Swe mice (age 3-6 months) [50]. Our analyses indicate no PA-induced changes to microvascular density, morphology, or surface area. For stimulus-induced hyperemic responses, the impacts of both Aβ and PA were most notably pronounced within the capillary bed, which accounts for the majority of vascular resistance and metabolite exchange [62]. In WT mice, PA yielded larger changes in stimulus- evoked capillary dilation and oxygenation changes; while in AD mice, PA yielded increased capillary dilation and faster, but smaller changes in capillary oxygenation.

Our observed differences in arterioles and capillaries support prior reports highlighting vulnerability of vascular smooth muscle cells, endothelial cells, and pericytes to Aβ, as well as the potential influences of PA on each of these cell types [60,61,80,80,82,83]. Importantly, while Aβ and PA alter their cellular phenotypes, our data reflects the resultant impact on vascular physiology.

The differential response of capillary oxygenation in WT and AD mice to forced PA from age 6-12 months constitutes a notable finding. In female C57BL/6N mice age 19-21 months, Shin et al observed that voluntary exercise yielded layer-specific increases in basal capillary pO2 and venous flow [84]. Conversely, we observed that capillary SO exhibited a transient decline following one month of routine, forced PA in male WT mice at age 6 months, which fully recovered by six months. This initial dip likely reflects a progressive physiological adaptation, where early increases in oxygen demand could outpace microvascular supply before to completely compensation with prolonged moderate exercise, a process associated with improvements in cerebral blood flow [85]. However, abrupt or excessive exercise may exceed the adaptive capacity of the cortical vasculature, potentially impairing the beneficial effects of moderate exercise [86]. This suggests that the neurovascular benefits of PA are contingent on duration, intensity, age, and possibly gender, emphasizing the importance of controlled exercise regimens in mitigating cerebrovascular deficits. Further studies are warranted to delineate the threshold at which exercise transitions from being beneficial to detrimental, particularly in the context of AD-related vascular dysfunction.

Our findings strongly support emerging hypotheses that emphasize the role of capillary dysfunction for AD progression [80,87–89]. Although it provoked no changes to capillary density or morphology in either WT or AD mice, routine engagement in aerobic PA shows great promise for strongly mitigating disturbances to basal capillary oxygenation and homogenize perfusion. Taken together, our basal pO2 measurements and functional hyperemia measurements offer insight to the “hyperactivation” phenomenon observed in fMRI BOLD studies of AD patients [90]. Although PA did not substantially affect absolute changes in SO2, the relative change is augmented by the reduced baseline in AD-noPA. When compared to other cohorts, our data indicates that reduced basal oxygen in the capillary and venous compartment account for a larger stimulus-evoked relative change.

While aerobic PA has proven useful for mitigating aging related alterations to vascular function, neuroinflammation, and numerous pathological hallmarks of preclinical AD [77,78,91–94], its protective mechanisms and capacity to prevent functional decline associated healthy aging and AD progression are still highly nuanced, multifaceted, and highly variable. Confounding parameters such as age, sex, intensity levels, and duration drastically affect PA’s effectiveness for neuroprotection. Our study highlights the benefits of maintaining routine aerobic PA throughout early adulthood and middle age to mitigate disruptions to cortical arteriolar and capillary oxygenation. Our findings motivate future investigations exploring PA’s effects on endothelial cells, vascular smooth muscle cells, and pericytes within cortical and subcortical brain regions.

## Materials and Methods

### Mice and Experimental Design

All animal handling and surgical procedures were in accordance with ARRIVE guidelines under a protocol approved by the Northeastern University Institutional Animal Care and Use Committee (IACUC). All male mice used in this study were on a C57BL/6 background. The APP/PS1dE9 transgenic mouse model (85Dbo/Mmjax, MMRRC Strain #034829-JAX, Jackson Laboratory) was utilized. Genotypes were confirmed via standard PCR reactions performed by Transnetyx Inc.. Mice were housed in a temperature- and humidity-controlled animal facility with a 14/10hour light/dark cycle and provided ad libitum access to food and water. Wild-type (WT) and Alzheimer’s disease (AD) mice were randomly assigned to sedentary (noPA) or physically active conditions (PA) (n=6/ cohort). All mice were bred and housed at the Northeastern University (NEU) vivarium, and all procedures were conducted in compliance with protocols approved by the NEU Institutional Animal Care and Use Committee (NE-IACUC).

### Treadmill Training

The experimental design was performed as previously illustrated in figure 1 [95]., previously assigned. A motor-driven treadmill (Ugo Basile, cat#47303) was used to train the animals at fixed speeds. To ensure recovery after cranial window surgery, treadmill training commenced 1-month post-surgery. Exercise mice underwent a 2-day acclimatization period, running for 10 minutes each day (Day 1 at 5 m/min; Day 2 at 8 m/min). Following acclimatization, exercise groups engaged in treadmill running at progressively increasing speeds (5–11 m/min) for 30 minutes per session, 5 days per week, for 5 months (from 6 to 12 months of age). Each session began at 5 m/min, with the speed incrementally increased to 11 m/min within the first minute. The maximum speed (11 m/min) was maintained for the remaining 29 minutes of the session. Mice in the sedentary groups were placed on the stationary treadmill for the same duration as the exercise groups but were not required to run. Animals that repeatedly failed to complete exercise sessions were excluded from the experiment. This long-duration training protocol has been reported to maintain exercise intensity at 45%–55% of VO max [96].

### Mapping the center of cortical activation

Cortical activation mapping was performed using a customized optical intrinsic signal imaging (OISI) system integrated into a commercial two-photon microscope (Ultima 2Pplus, Bruker) equipped with a high-resolution monochromatic camera (acA1300-200μm, Part No. 106752, Basler aceU). A continuous broadband LED light source (KL1600 LED, Part No. 1500-600, Schott), filtered at the isosbestic wavelength of oxy- and deoxyhemoglobin (bandpass filter, 568/4 nm, cat# 65160, Edmund Optics), provided illumination. Hemodynamic responses were recorded at 5 Hz [97] to capture stimulus-evoked cortical activity. Activation ratio maps were generated by normalizing the post-stimulus signal (0–1.5 s after stimulus onset) to the baseline period (1–4 s prior to stimulus onset). The spatial localization of cortical activation (300 × 300 µm) was determined using a custom algorithm that mapped the entire cranial window. After applying the 3D median and 3D Gaussian filters to the image series, the activation center was determined by averaging the vectorized output of the region of interest (ROI) across successive imaging frames.

### Chronical Cranial Window Implantation

Surgical procedures for implantation of chronic cranial windows were performed as previously described [98]. Briefly, mice were pre-treated with dexamethasone (4.8 mg/kg) and cefazolin (0.5 g/kg) 4–6 hours before surgery to prevent brain edema and infection, respectively. Mice were anesthetized with isoflurane (2% for induction and 1%–2% during surgery). A circular incision was made on the scalp over the frontal and sagittal bones, and the exposed periosteum was carefully removed with a No. 15 blade to ensure proper fixation. A custom holding bar was attached to the skull using cyanoacrylate adhesive (Loctite-401, Henkel, Cat#135429) to immobilize the animal during imaging. A craniotomy (∼3 mm in diameter) was performed on the left hemisphere over the primary somatosensory cortex (coordinates: A–P: 2.0 mm; M-L: 3.0 mm) while leaving the dura intact. The craniotomy was sealed with a glass plug and secured with dental cement. Post-surgical care included administering Sulfamethoxazole/Trimethoprim (0.5 mg/0.1 mg/mL) and Carboprofen (1 mg/mL) in drinking water for 5 days. Additionally, cefazolin (0.5 g/kg) was given via intraperitoneal injections twice daily for 5 days, and Buprenorphine (0.05 mg/kg) was administered subcutaneously once daily for the first 3 days to manage pain. Mice were provided with softened food pellets immersed in water in a petri dish placed on the cage floor to ensure adequate nutrition and hydration during recovery.

### Training for awake imaging

Two weeks after surgery, to ensure full recovery, mice were trained to tolerate head fixation in a custom-designed cradle for imaging sessions lasting up to 2 hours. Training sessions were conducted three times per week over a 2-week period, with the duration of each session gradually increased. During training and imaging sessions, mice were rewarded with sweetened milk approximately every 15 minutes. To promote comfort and natural behavior, mice were placed on a suspended fabric bed, allowing them to engage in natural grooming behaviors

### Functional activation protocol

Functional hyperemia in the somatosensory primary cortex (Barrel cortex) was induced by pneumatic whisker deflection, following a previously established protocol [99]. Briefly, a Picospritzer microinjection device (PDES-01DXH-E-LA-4, npi electronic GmbH). Pneumatic air puffs (10–20 psi) were delivered to the contralateral whisker pad relative to the cranial window implantation. To ensure effective activation of the whisker Barrel cortex while minimizing the eye-blink reflex, air puffs were applied in an inferoforward direction. A 3D-printed custom nozzle directed the air puffs toward the E1 whisker. As other whiskers were not trimmed, the stimulus affected multiple whiskers simultaneously. Each trial consisted of a 5-second baseline period followed by a 1.5-second stimulation period. During stimulation, air puffs were delivered at a frequency of 3.3 Hz, with each puff lasting 300 ms and separated by a 300-ms interval. The inter-stimulation interval (ISI, onset-to-onset) was set to 35 seconds.

### Oxygen Measurements

Systemic oxygen delivery and exchange were quantified using OxyPhor-2P, a phosphorescent oxygen-sensitive dye injected intravascularly (10 μL of 20 μM solution). Two-photon excitation fluorescence microscopy was conducted with a scanning microscope (Ultima 2Pplus, Bruker), integrated with a tunable ultrafast laser (80 MHz, <120 ps pulse width, Insight X3+, Spectra-Physic) tuned to 950 nm and a 25× water-immersion objective (380–1050 nm, 1.1 NA, 2.0 mm WD, N25X-APO-LWD, Nikon). System control, including motorized alignment with the cortical surface, image acquisition, and adjustments to laser power and PMT voltage, was managed using commercial software (Prairie View, Bruker), fully integrated with the scan head, laser power and PMT voltage control.

Measurements of partial pressure of oxygen (pO2) were derived from the phosphorescent lifetime of OxyPhor-2, utilizing the Stern-Volmer equation as previously described [100]. Each phosphorescence excitation/collection cycle comprised a 10 μs excitation pulse train (800 femtosecond pulses) followed by a 290 μs interval with no laser excitation to collect phosphorescence decays, culminating in a total cycle duration of 300 μs. At each cortical depth, data acquisition was conducted across 10 microvessels, with 100 excitation/collection cycles per timepoint. This process was riterated to measure ∼20 seconds pO2 transients per stimulus trial, for a total of 18 stimulus trials per cortical depth. Emission light was routed through an infrared blocker (short-pass filter, 900 nm) and a dichroic mirror (bins splitter, 565 nm) into a four-channel detection system. The phosphorescence emission was isolated with a 795±75 nm bandpass filter and detected via a GaAsP photomultiplier tube (PMT; H10770PA-50/001, Hamamatsu). Emission signal was counted via time-correlated single photon counting module (SPC-150N TCSPC, Becker & Hilks). Phosphorescence lifetimes were determined using custom software by fitting decay curves to single-exponential models using non-linear least squares regression. Standard errors for lifetime measurements were estimated through a bootstrap resampling methodology [100]. The calculated lifetimes and associated standard errors were converted to pO2 values using a fitted calibration profile. Oxygen saturation (SO2) was calculated from pO2 measurements using the Hill equation, with a Hill coefficient (h) of 2.59 and a P50 value of 40.2 mmHg, as previously described [99].

### Capillary blood flow imaging and vascular dynamic with two-photon laser-scanning fluorescence microscopy

Capillary flow dynamics were quantified using the Ultima2pPlus microscope as previously detailed (refer to Oxygen Measurement section). Vascular labeling was achieved through retroorbital injection of 10 μL FITC–Dextran 70 (50 mg/mL in saline; Sigma-Aldrich, cat# 46945). This fluorescent marker delineated the plasma as bright regions in the images, with red blood cells (RBCs) appearing as dark shadows. Imaging was performed within a selected region of interest (600 μm × 600 μm) defining by OISI protocol, targeting approximately 10 capillaries (range: 7–13 capillaries; diameter <10 μm) across a cortical depth range of 0–400 μm, with measurements obtained at 50 μm depth intervals. Velocity calculations involved tracing a straight line parallel to the longitudinal axis of each capillary, while internal diameter measurements were obtained by drawing perpendicular straight line across the lumen at approximately 50 points of length each one. Data acquisition was conducted continuously during a functional activation protocol. Velocity computation was performed using a MATLAB-based dedicated function: HydroVel [101], and lumen diameter analysis was facilitated by a custom image processing routine. Acquired data were aggregated into 1/100-second time bins for subsequent analyses.

### Microvascular Morphology Measurements

Microvascular angiograms were recorded within the same regions of interest (ROIs) used for functional measurements (e.g., intravascular pO2, blood flow, and arteriolar diameter transients). Imaging was performed on a separate day in awake mice at rest. Three-dimensional fluorescence intensity stacks were acquired down to a depth of 400 μm from the cortical surface.

### Data Analysis

Oxygen saturation levels were quantified in arterioles, capillaries, and venules, alongside measurements of blood flow velocity, vascular dynamics, and vessel density, across defined cortical layers. To assess vascular heterogeneity under both resting and stimulus-evoked conditions, the coefficient of variation (CV) of microvascular oxygen tension was calculated within 50-µm cortical depth bins for each individual animal. Statistical comparisons across experimental groups were performed using linear mixed-effects models implemented with the lme4 package (version 1.1.37) in R (version 4.4.2, Pile of Leaves). The models included group and vascular segment as fixed effects and animal identity as a random intercept, accounting for repeated measures within subjects. Post hoc pairwise comparisons were conducted using estimated marginal means with Tukey adjustment for multiple comparisons, implemented via the emmeans package (version 1.11.0). All results are reported as mean ± standard error of the mean (SEM), and statistical significance was defined as *p*: *< 0.05, **<0.01 and ***<0.001.

## ACKNOWLEDGMENTS

This work was performed with generous support from the Northeastern College of Engineering and the National Institutes of Health: NIH R01AA27097, NIH R56AG058849, and R21AG085655. We thank Dr. Sava Sakadzic and Allen Alfadhel from the MGH Martinos Center for Biomedical Imaging for consultations with data processing and interpretation. We thank the Institute for Chemical Imaging of Living Systems (RRID:SCR_022681) at Northeastern University for consultation and imaging support, and for the assistance of confocal imaging and wide-field fluorescence imaging.

