## Supplementary Figure for "Chronic Aerobic Exercise Alleviates Amyloid-induced Capillary Dysfunction"

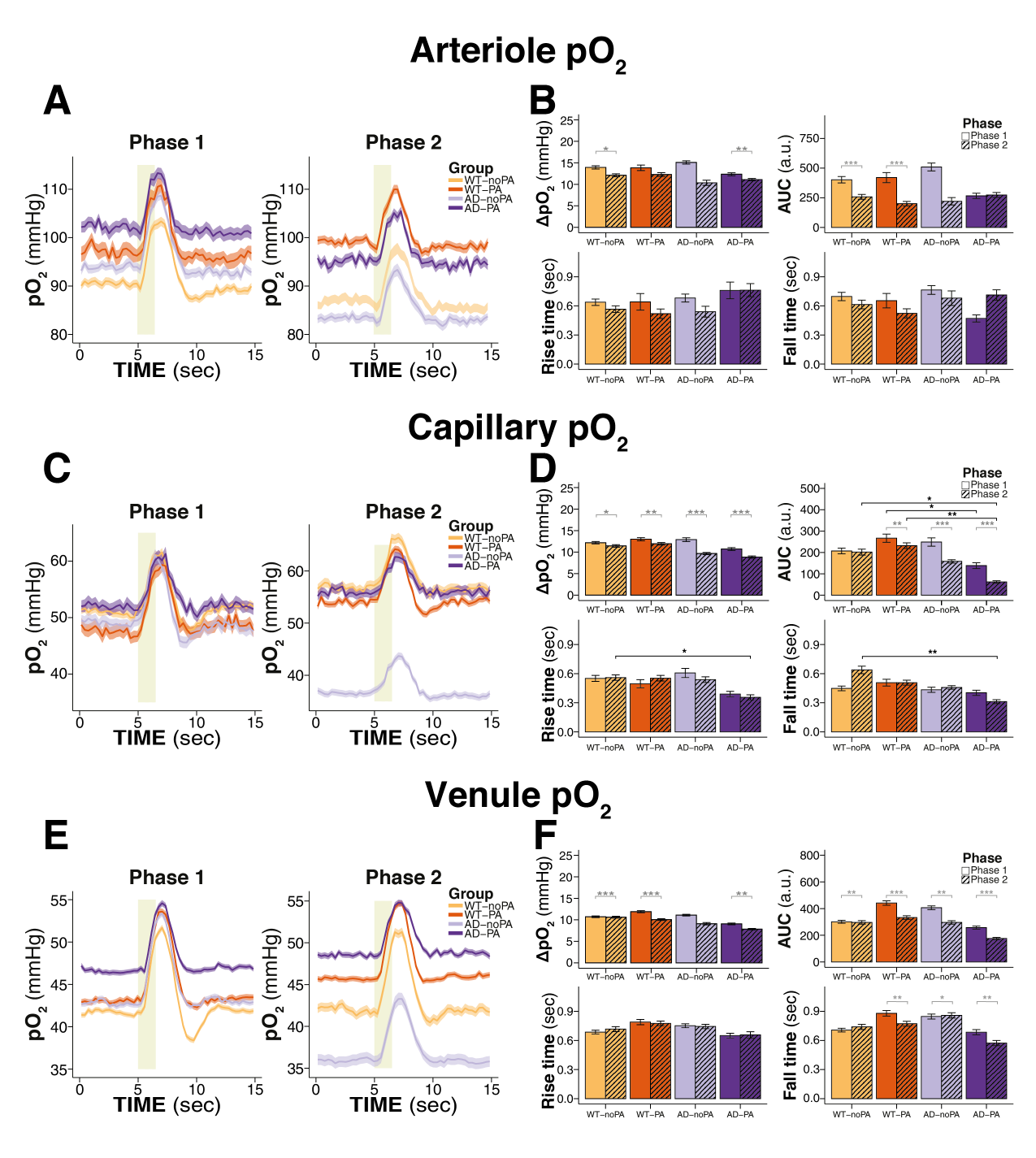

**Figure S1** pO_2_ transients during whisker stimulation in wild-type and AD mice under aging and physical activity (complementing the SO_2_ transient data from manuscript figure 4): *(A-B)* Arteriolar oxygen transients show higher basal pO_2_ *(C-D)* In capillary vessels, Aβ and aging yielded reductions in basal pO_2_ in AD-noPA mice. Routine PA protected AD-PA mice from reductions in basal pO_2_. In WT animals, hyperemic increases in pO_2_ were higher in PA animals compared to sedentary noPA animals, while in AD animals, PA animals responded to stimulation with smaller, yet faster changes in pO_2_. (E-F) In venules, differences in basal pO_2_ were higher in WT-PA and AD-PA animals at phase 2. Functional stimulation evoked smaller venous SO_2_ changes at phase 2 than phase 1.

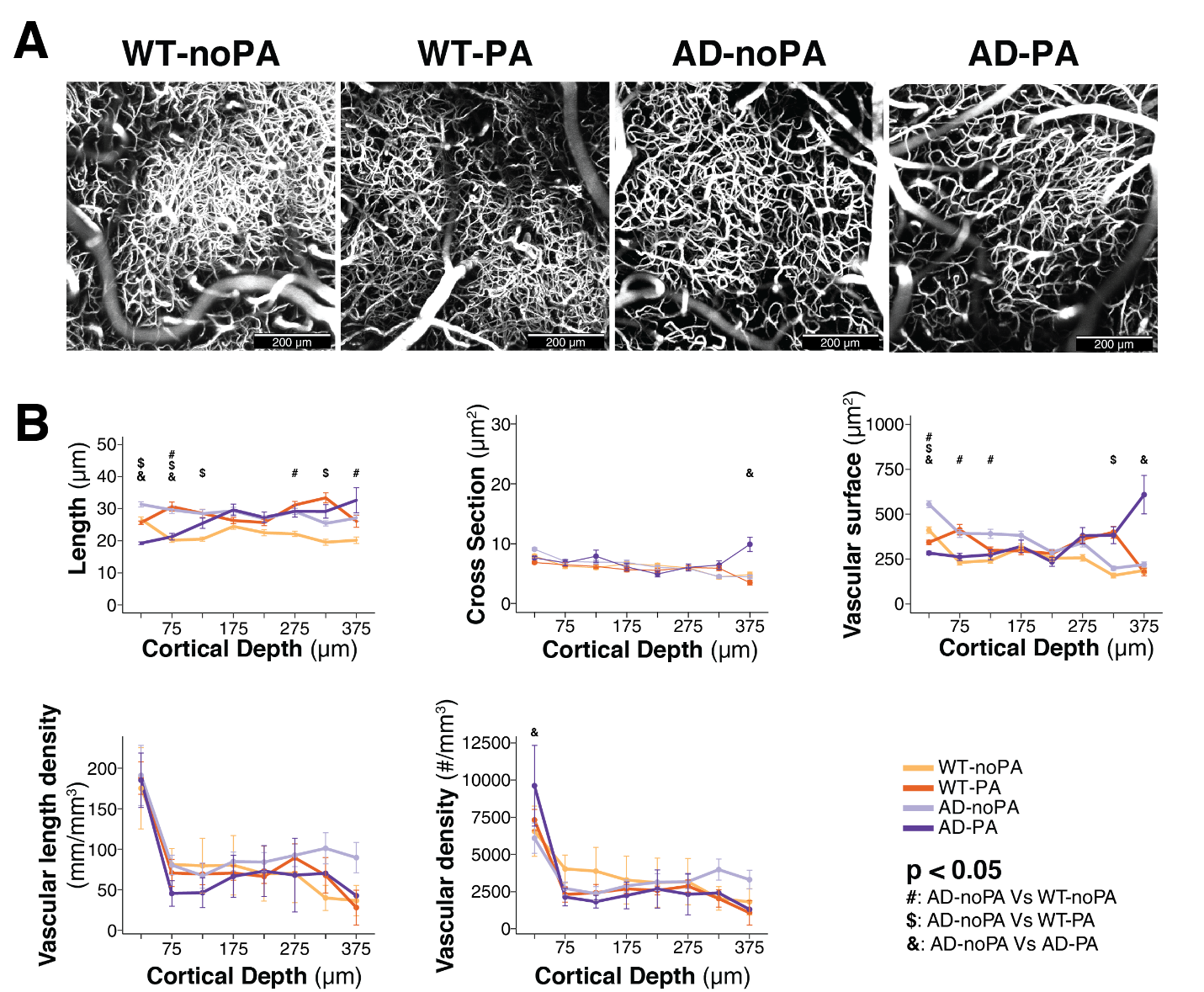

**Figure S2** Comparison of microvascular architecture from 4 experimental cohorts at Phase 2, segmented and analyzed with Matlab-based VIDA software package . (a) representative MIP images of vascular stacks from somatosensory cortex in WT-noPA, WT-PA, AD-noPA, and AD-PA cohorts. (B) Mean vessel length, cross sectional area, vascular surface area, vascular length density, and vascular density, all plotted against cortical surface depth. 6 months of routine PA did not yield substantial variations of these metrics in AD or WT mice.
